# Study the dynamics of behavioral and biochemical parameters in the *PARK2*-knock-out mice model of Parkinson’s disease

**DOI:** 10.64898/2026.03.02.709182

**Authors:** Ekaterina A. Emelianova, Olga A. Averina, Oleg A. Permyakov, Anastasia V. Priymak, Mariia A. Emelianova, Olga O. Grigoryeva, Svetlana A. Garmash, Petr V. Sergiev, Olga U. Frolova, Kirill E. Kianitsa, Vladislav S. Savitskiy, Maxim L. Lovat

**Author notes:** The author responsible for the correspondence: Ekaterina Emelianova; 1451 East 12^th^ Ave, Vancouver, BC V5N 0J9.

## Abstract

**Background:** Parkinson’s disease (PD) is a progressive chronic neurodegenerative disease. The *PARK2* gene encoding the Parkin protein accounts for approximately half of early-onset autosomal recessive PD cases in humans.

**Objective:** The aim of this work was to study the effect of the *PARK2* gene knockout in mice on the dynamics of behavioral and biochemical parameters of PD.

**Methods:** The study was performed on C57BL/6-line mice aged from 4 months to 1.5 years: wild type (*park2 +/+*), heterozygotes (*park2 +/-*) and homozygotes (*park2 -/-*) knocked out by the *PARK2* using CRISPR-Cas9. The open field test, the Porsolt forced swimming test, the grid-walk test, the beam-walking test, the elevated plus maze test, the accelerating rotarod test were used to assess the behavioral phenotype. Measurement of the concentration of bioamines and their metabolites by HPLC and evaluation of the amount of tyrosine hydroxylase, BDNF and GDNF by Western Blot were used to study the biochemical signs of PD.

**Results:** *Park2 -/-* mice begin to show signs of decreased motor activity no earlier than at 4 months of life. At 12 months of life, it was shown only a decrease in the level of the mature isoform of GDNF and an increase in the number of immature isoforms in the frontal cortex and striatum were revealed.

**Conclusion:** The data obtained indicates a different age dynamic of the condition of mice associated with the *PARK2* knockout. However, no pronounced specific manifestations of PD in human were found in *park2 -/-* mice.

## Introduction

Parkinson’s disease (PD) is a slowly progressive chronic neurodegenerative disease characteristic of the older age group. It is the second most common neurodegenerative disease, which affects about 2% of the population over the age of 60.^1^

This disorder is characterized by progressive degeneration and death of neurons that produce dopamine in the substantia nigra and other subcortical nuclei responsible for motor functions. Insufficient dopamine production leads to an inhibitory effect of the basal ganglia on the cerebral cortex, due to which their ability to initiate and smooth out voluntary muscle movements is suppressed. The etiology of the disease has not been studied enough. The main risk factors are genetic predisposition, aging and/or environmental factors. Mutations in several genes are known to cause autosomal dominant and recessive forms of PD. In particular, mutations in the *PARK2, SNCA*, and *PINK1* genes may be the cause of familial forms of PD, whereas various genetic defects at other loci may be associated with sporadic forms of PD without a family history.^2,3^ It is known that the *PARK2* gene encoding the Parkin protein is responsible for half of the cases of early autosomal recessive PD in humans.^4^

The Parkin is an E3-ubiquitin ligase and is involved in maintaining protein and mitochondrial homeostasis, namely, in proteolysis of damaged proteins, autophagy of mitochondria, and also performs a direct antioxidant function.^5^ This enzyme is expressed mainly in tissues whose functioning is associated with high energy consumption, such as the heart, muscles, retina, and brain.^6^ Catecholaminergic neurons are particularly sensitive to damage, since catabolism of catecholamines is accompanied by direct formation of peroxide and their synthesis is associated with high activity of the mitochondrial respiratory chain.

Consequently, mutations in the *PARK2* gene lead to impaired functioning of the Parkin protein, accumulation of damaged proteins, mitochondrial dysfunction, and neuronal death (primarily in catecholaminergic neurons, as the most energetically loaded).^7^

There are a lot of neurotoxic *in vivo* models of PD mostly aimed at inhibiting complex I of the respiratory chain, which leads to oxidative damage of neurons.^8^ Currently increasing attention is being paid to genetic models of PD, since one of the risk factors for disease is a genetic predisposition. The most used knockout genes among them are *PARK2* (Parkin), *PARK6* (PINK1), and *PARK7* (DJ-1). Many animal models of PD cannot simulate the progressive and protracted nature of the disease, which may be crucial for the correct definition of neuroprotective therapy. In addition, there is no animal model capable of reproducing all the signs of idiopathic Parkinson’s disease in humans. Changes in the content of tyrosine hydroxylase (TH) and other dopamine markers are not always observed in transgenic mouse models, while α-synuclein aggregation is rarely observed in toxic models. Cognitive impairment is rare, and if it occurs, it is usually early and is not associated with the typical histopathology of dementia in Parkinson’s disease.^9^

Accordingly, the aim of this work was to study the effect of knockout of the *PARK2* gene in mice of different ages on the behavioral and biochemical parameters of Parkinson’s disease.

## Materials and methods

The study was performed on male and female mice of the initial C57BL/6-line SPF category (specific pathogen free) weighing 25-30 g and aged from 4 months to 1.5 years. The animals were kept in standard conditions (temperature 22 ± 1 ° C, humidity 40-70%, light cycle - 12 hours) with unlimited access to non-pyrogenic water and autoclaved standard feed. The mice were divided into 3 groups: wild type (*park2* +/+), heterozygotes (*park2* +/-) and homozygotes (*park2* -/-) knocked out by the *PARK2* gene using CRISPR-Cas9 technology (reading frame shift mutation in exon 5 of the *PARK2* gene). Mice C57Bl/6J and CBA (Federal Research Center Institute of Cytology and Genetics, Siberian Branch, Russian Academy of Sciences (ICG SB RAS) (Novosibirsk, Russia), SPF status, were mated to obtain F1 hybrids C57Bl/6J × CBA. Zygotes were obtained by mating of the hybrid super ovulated female mice with males using standard procedure.^10^ Mice were genotyped for *PARK2* gene. The distribution of animals into groups (n) is indicated below. Experiments were conducted in accordance with Directive 2010/63/EU of the European Parliament and the Council of the European Union. The protocol of the experiment was approved by the Bioethics Commission of the Lomonosov Moscow State University Mitoengineering Research Institute. Protocol No. 142 dated 09.09.2019.

## 1. Investigation of behavior parameters

### 1.1. The study of motor activity in the open field test

A modification of the open field test was used in this study: a square arena with a side of 24 cm (RPC OpenScience Ltd, Russia) illuminated by bright light (500 Lux).^11^ The following parameters were recorded using the “Ethovision XT 14” program: distance (in cm), movement time (at a speed of more than 5 cm/sec), immobility time (at a speed of less than 0.2 cm/sec), average speed (cm/s), the number of episodes of mobility and immobility. The same set of parameters, as well as the latency period and duration of stay, were recorded for the central sector. The total registration time for each animal was 5 minutes.^12^

### 1.2. The study of the activity of mice in the forced swimming test

The test was used to assess general motor activity, as well as behavior strategies in a state of acute non-runaway stress.^13^ The mouse was placed for 10 minutes in a transparent vessel 30 cm high, 10 cm in diameter, filled with water (temperature of water +21-23 °C) to a mark at a height of 25 cm. The duration and number of motor acts were recorded for active (vigorous movements with all paws), passive (weak strokes with 1-3 paws) swimming, as well as immobility (immobilization).^14^ The mice were placed in a heated cage with napkins until they dried after the procedure. The behavior indicators in this test were processed using the “Realtimer” program (RPC OpenScience Ltd, Russia), which allows us to record the sequence of events, their duration, and number, as well as conducting basic statistical data processing.

### 1.3. Assessment of motor deficits and neurological status of mice in the grid-walk test

Motor deficiency was assessed by the relative number of errors committed by animals during free movement for 5 minutes on the surface of a horizontally installed grid measuring 25*35 cm, with a bar diameter of 1 mm, with a cell of 1 cm. The number of cases of free slipping of mouse paws into cells – n (motor errors), as well as the total number of steps taken (an indicator of the total motor activity of the animal) - n_(total)_ were recorded with the help of a side–mounted DVR. The results were presented in the form of a normalized number of motor errors per 100 animal steps (n/n_(total)_*100).^15,16^ The mice underwent a 15-minute adaptation in a home cage to obtain more reproducible data, which made it possible to neutralize the stress factor in the experiment.

### 1.4. Assessment of the neurological status of mice in the beam-walking test

The test consists of a horizontal tapering step bar with a length of 100 cm, fixed at a height of 100 cm, with a shelter at its narrow end (RPC OpenScience Ltd, Russia) and a side light source of 500 Lux. The animal walks along the bar, which gradually narrows, and enter a dark compartment (shelter). The tapering beam has a lower stand (a transparent beam, 1 cm wider than the upper one). The width of the upper beam narrows from 2.5 cm to 1 cm. The movement of the animal is recorded on the camera in high resolution. There is a mirror on the opposite side of the camera to assess the movement of the paws on both sides of the animal.^17^

The number of limb positions on the lower bar (errors), the number of slips from the upper bar to the lower bar (when the front or rear paw is placed on both bars), and the total number of steps taken from the starting line to the animal entering the dark compartment are calculated, for each front and rear paw separately. The data obtained for the three attempts is averaged. The degree of severity of sensorimotor deficiency (SD) is calculated using the formula as a percentage:

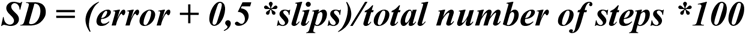

### 1.5. The study of mice in the elevated plus maze test

The test is designed to assess the level of locomotor activity, as well as the balance between research activity and anxiety.^18^ The installation consists of 2 closed (illumination 50-70 Lux) and 2 open (illumination 700 Lux) beams located opposite each other (beam length 30 cm). The height of the sides of the closed beams is 15 cm. The installation is raised 70 cm above the floor level. The animal is placed in the center of the maze with its head facing the open beam. The program "Noldus Ethovision XT 14" automatically registers within 5 minutes the following parameters: distance (in cm), time of movement (at a speed of more than 2 cm/sec), time of immobility (at a speed of less than 0.2 cm/sec), average speed, number of episodes of mobility and immobility. The same set of parameters, as well as the latency period and duration of stay, is fixed for the central sector, open, and closed beams separately. The animal Anxiety Index (AI) is calculated using the formula: AI=100*(1-(Tl/Ot + Nl/Nt)/2), where Tl is the time in the light, Ot is the total testing time, Nl is the number of transitions to the light, Np is the total number of transitions between the installation beams.

### 1.6. The study of mice in the accelerating rotarod

The installation is a rotating steel rod (45 cm in length, 4 cm in diameter) covered with non-slip rubber (LLC “Neurobotics”, Russia). The rod is divided into eight compartments (4 cm) by means of plastic discs. The rotation speed of the drum increases smoothly from 3 to 30 revolutions per minute for 5 minutes. The test recorded the parameters measured for 6 consecutive attempts by animals to stay on the installation: the maximum holding time on the rod and the number of attempts at which this maximum was reached. The test allows to evaluate motor disorders, endurance, formation of motor skills.

## 2. The study of biochemical parameters

Cervical dislocation of animals was performed at the 12th month of life with sampling of frontal cortex and striatum tissues.

### 2.1. Measurement of the concentration of major bioamines and their metabolites by HPLC

Solutions of norepinephrine (NE), dopamine (DA), serotonin (5HT), dioxyphenylacetic acid (DOPAC), homovanillic acid (HVA), 5-hydroxyindolacetic acid (5HIAA) at a concentration of 50 ng/ml for the frontal cortex and striatum were used as external standards.

The concentration analysis was performed on an Eicom HTEC-500 chromatograph (Japan). The C-18 column CA-50DS was used. The composition of the mobile phase was: 0.1 M phosphate buffer, EDTA, sodium octane sulfate, 18% methanol, pH =4.5. The potential at the electrode was 650 mV; the flow rate of the mobile phase was 200 µl/min.

The peak areas were compared to the peak areas of external standards, which were measured every 10 samples, to determine the concentration. Rationing was also carried out for the peak area of the internal standard.

### 2.2. Evaluation of the amount of TH, BDNF and GDNF by Western blotting

Protein extraction was performed after sampling of the frontal cortex and striatum using a TRI-reagent. The proteins in the samples were separated by SDS-PAGE in 10% PAAG. Protein fractions from the gel were transferred to the PVDF membrane. According to protein standards, a section corresponding to the molecular weight of the protein of interest was cut out of the membrane (∼55 kDa for TH; ∼10-25 kDa for BDNF; ∼15-35 kDa for GDNF). Incubation was carried out in a solution of primary (anti-TH, ab112, Abcam, rabbit, 1:1000; anti-BDNF, ab108319, Abcam, rabbit, 1:5000; anti-GDNF, ab18956, Abcam, rabbit, 1:1000) and secondary antibodies (anti-rabbit HRP conj., 1:2000 for TH; 1:5000 for BDNF; 1:1000 for GDNF). Signal strength was visualized and evaluated using the ChemiDoc MP (BIO-RAD) system.

## 4. Statistical data analysis

The analysis was performed using the GraphPadPrism8 program (GraphPadSoftwareInc., USA) using the Student’s t-test, one-way analysis of variance (One-way ANOVA), two-way analysis of variance (2way ANOVA). The data was expressed as median ± interquartile range. The differences at p <0.05 were considered statistically significant (*/#).

## Results and discussion

1. The age dynamics of the behavior of mice knocked out by the *PARK2* gene.

It is known that PD is accompanied by the gradual development of motor activity disorders in elderly patients. Therefore, we studied the age dynamics (from 4 to 18 months) of motor activity of mice knocked out by the *PARK2* gene.

Heterozygotes did not differ from the wild type in any of the behavior parameters. Mice aged 4 months *park2* -/- in the open field test demonstrated a statistically significant decreased distance compared to the wild type (data is not presented). It was also found that *park2* -/- males achieve maximum holding time on later attempts than control mice, although without statistically significant differences in maximum retention time in the accelerating rotarod test (Fig.1, a). The data obtained may indicate both a lower reaction to a negative stimulus (the risk of falling off the rod) and a decrease in the rate of motor skill formation.

**Fig 1.**
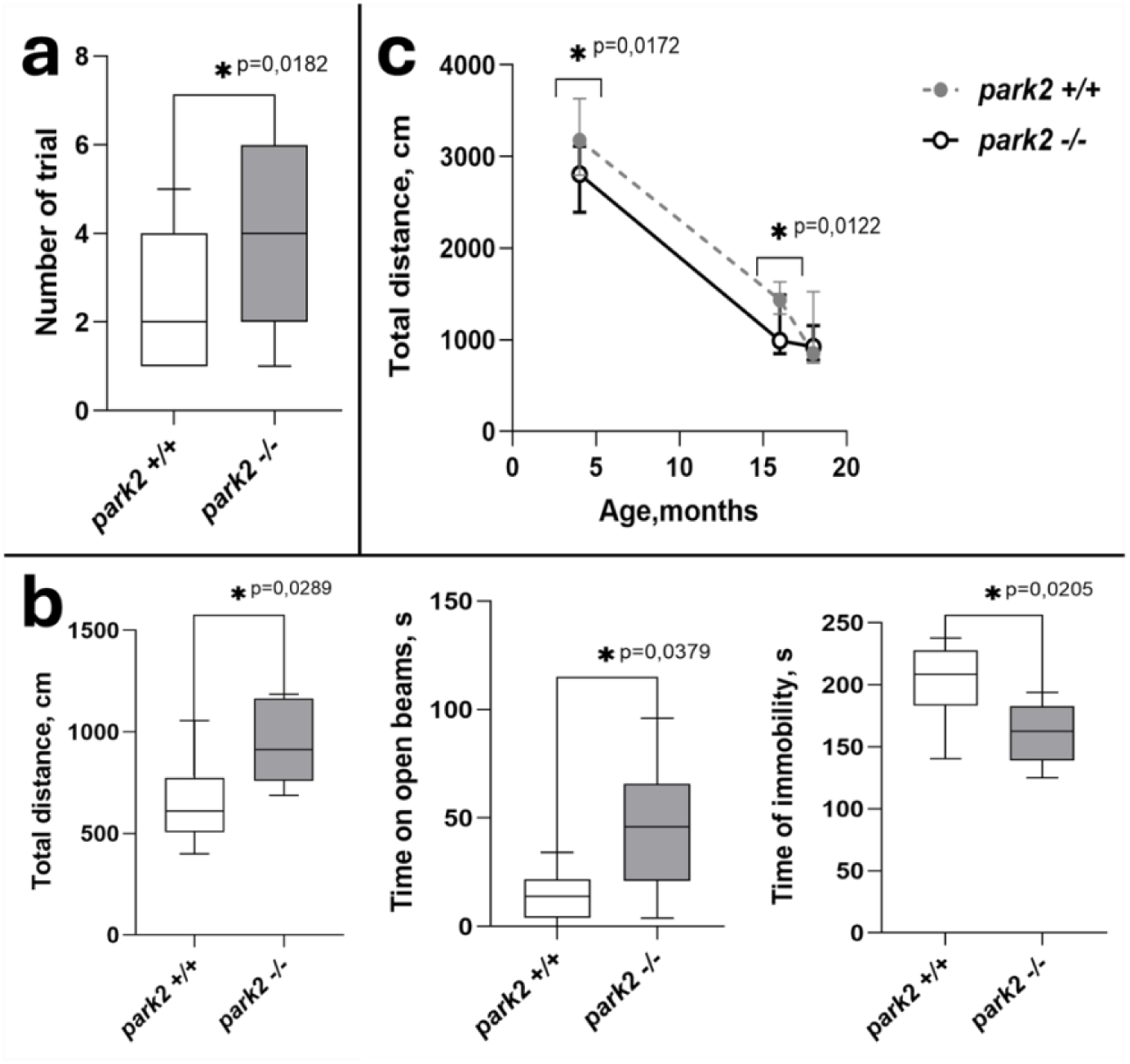
Comparison of the behavior of 4-month male mice **(a)** park2 +/+ (n=16) with park2 -/- (n=15) in the accelerating rotarod test; **(b)** park2 +/+ (n=8) with park2 -/- (n=7) in the elevated plus maze test; **(c)** 4-months park2 +/+ (n=11), park2 -/- (n=24); 16- months park2 +/+ (n=12), park2 -/- (n=11); 18-months park2 +/+ (n=5), park2 -/- (n=15) in the open field test.

Knockouts at the age of 4 months had a shorter immobility time with a longer distance covered and time spent on the open beams of the elevated plus maze test (Fig.1, b). The effects changed at a more mature age (starting from 18 months of life). Differences in distance in the open field test disappeared due to a faster age decrease in activity in the control group (Fig.1, c). Moreover, knockouts stay longer in the center of the open field starting from 16 months age compared to wild-type mice (Fig.2, a).

**Fig 2.**
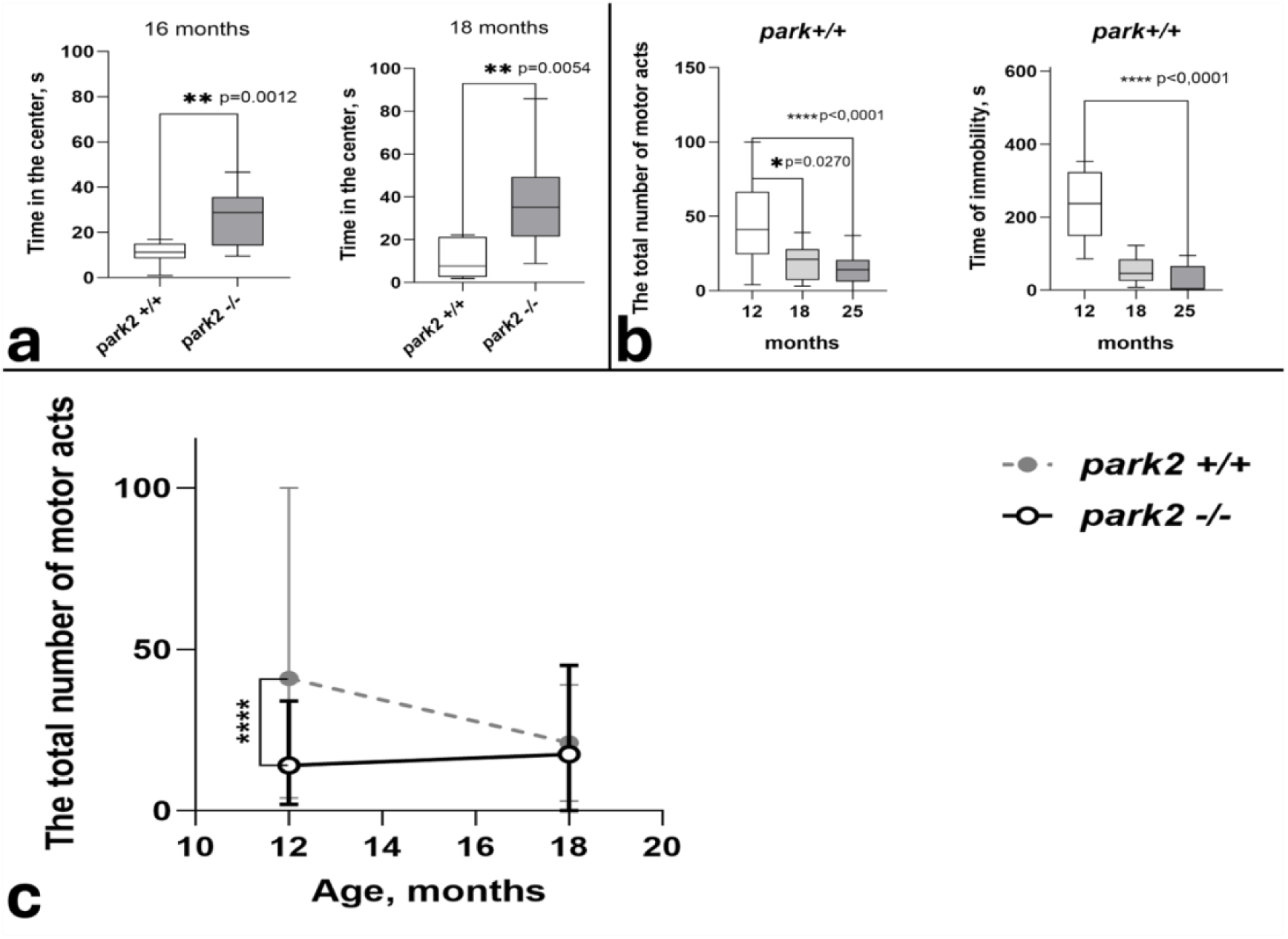
**(a)** Comparison of the behavior of male mice (16-months park2 +/+ (n=12), park2 -/- (n=11); 18-months park2 +/+ (n=5), park2 -/- (n=15)) in the open field test. Mann Whitney test. **(b)** Age dynamics of the immobility time and the total number of motor acts in male mice park2 +/+ 12-months (n=10); 18-months (n=5) and 25-months (n=12) in the Porsolt forced swimming test. One-way ANOVA, Dunn’s multiple comparisons test. **(c)** Age dynamics of the total number of motor acts in the Porsolt forced swimming test of male mice park2 +/+ (12-months, n=30; 18-months, n=50) and park2 -/- (12-months, n=25; 18-months, n=15). Unpaid t test. The data is presented as a median, the spread is shown as an interquartile range.

The dynamics of animal activity in the forced swimming test in the control group shows a decreasing immobility time with age. Statistically significant differences developed by 25 months of animal life (Fig.2, b). These results may reflect the predominance of an active defensive reaction of animals to stress and a decrease in depression, which is consistent with the literature data.^19^

The total number of motor acts decreased with age reflecting the oppression of the behavioral repertoire (Fig.2, b). The age dynamics of the behavior of <u>knocked out</u> males and females were different. Thus, the number of motor acts in this group of animals was low already at the age of 12 months (Fig.2, c) with a similar dynamics of decreasing immobilization time (data is not presented). The data obtained is consistent with the results of the open field, the accelerating rotarod test and reflect the suppressed reaction to stressful environmental conditions in the knocked-out animals.

Evaluation of the dynamics of neurological and motor status in knockouts in the grid-walk test showed a statistically significant increase in the number of motor errors with age in both knockout and wild-type mice. Unlike previous tests, the dynamics of the sensorimotor status of the knockouts in the beam-walking test did not differ (graphs are not shown).

Thus, a decrease in the depressive-like state is observed with an increase in the number of motor errors in adult control C57BL/6 mice. At the same time, mice knocked out by the *PARK2* gene are less anxious in adulthood. A slight decrease in general motor activity observed is leveled with age. That is, knockout of the *PARK2* gene in mice does not lead to accelerated motor dysfunction and depression even by the age of 25 months (close to the limit for these animals). Opposite, it is known that mutations in the *PARK2* gene in humans are accompanied by a risk of developing PD with an early onset and a characteristic dynamics of decreased motor activity.

A statistically significant decrease in motor activity, the development of a depressive-like condition, and an aggravation of sensorimotor deficiency were shown in our previous studies on mice, where pharmacological methods (administration of protoxin MPTP) were used to simulate PD and a similar set of behavioral tests were implemented, which also does not coincide with the development of disorders in *PARK2* knockout mice.^20^

It has been shown in recent work on sequencing the human and mouse genomes that to date only about 300 genes out of 20,000 are presumably unique and different in these taxa.^21^ However, despite the high genetic homology and conservativeness, even structurally similar human and mouse proteins do not always perform identical functions. Some evidence indicates that the genetic deletion of the *PARK2* gene in mice causes a violation of the synaptic plasticity of the hippocampus, which leads to impaired memory without serious violations of olfaction or motor skills.^22^ Another group demonstrated that *park2* -/- mice display impaired habituation to a new environment and exhibit increased thigmotaxic behavior and anxiety-related parameters in the light/dark test that may reflect anxiety disorders in PD.^23^ Our results support the evidence that Parkin protein functions in mice may differ from those in humans. Moreover, it was found that there are some compensatory mechanisms that might mitigate Parkin deletion in mice. So, it was shown that Syt11, a potent substrate of Parkin, exhibited increased expression during the suckling stage in Parkin KO mice but returned to normal levels in adulthood, which might underscore the role of compensatory Syt11 expression during development in the absence of PD phenotypes in Parkin KO mice.^24^

2. Assessment of the content of basic bioamines and their metabolites in the striatum.

The morphological basis of PD is the death of dopaminergic neurons, which is accompanied by a decrease in the level of dopamine and its metabolites in the brain. Moreover, it has been shown that there is also a decrease in the level of other bioamines in patients with PD. Therefore, we measured the concentration of the main bioamines by HPLC (Table 1).

**Table 1.** The content of basic bioamines and their metabolites in the striatum.

| Concentration of bioamines, nmol/g | Genotype |  |
| --- | --- | --- |
|  | *park2 +/+* | *park2 -/-* |
| **DA** | **40.77 ±26.08** | **37.22 ±23.6** |
| **DOPAC** | **1.840 ±1.205** | **1.400 ±0.666** |
| **[DOPAC/DA]** | **0.055 ±0.026** | **0.048 ±0.025** |
| **5-HIAA** | **8.840 ±1.893** | **8.817 ±2.276** |
| **HVA** | **2.690 ±1.240** | **2.217 ±0.624** |
| **3-MT** | **1.770 ±1.148** | **1.683 ±0.697** |
| **5-HT** | **7.690 ±2.455** | **7.233 ±1.513** |

| Concentration of<br>bioamines, nmol/g | Genotype |  |
| --- | --- | --- |
|  | *park2* +/+ | *park2* -/- |
| **DA** | **40.77 ±26.08** | **37.22 ±23.6** |
| **DOPAC** | **1.840 ±1.205** | **1.400 ±0.666** |
| **[DOPAC/DA]** | **0.055 ±0.026** | **0.048 ±0.025** |
| **5-HIAA** | **8.840 ±1.893** | **8.817 ±2.276** |
| **HVA** | **2.690 ±1.240** | **2.217 ±0.624** |
| **3-MT** | **1.770 ±1.148** | **1.683 ±0.697** |
| **5-HT** | **7.690 ±2.455** | **7.233 ±1.513** |

In addition to the total content of dopamine and its metabolites, dopamine turnover was also calculated from the ratio [DOPAC]/[Dopamine]. An increase in this coefficient reflects the predominance of degradation of this mediator over its synthesis and can be used as a marker of the progression of neurodegenerative disorders.

An assessment of the content of basic bioamines and their metabolites in the striatum of all the groups at the age of 12 months showed no statistically significant differences between them. The turnover of bioamines also did not differ.

3. Assessment of the content of TH and neurotrophic factors in the brain of mice.

The concentration of TH in the striatum and in the frontal cortex of mice at the age of 12 months measured by Western blotting (Fig.3, a). does not differ. The data obtained is consistent with the HPLC results. Probably, knockout of the *PARK2* gene in mice at the age of 12 months does not affect the synthesis and metabolism of bioamines in the brain.

**Fig 3.**
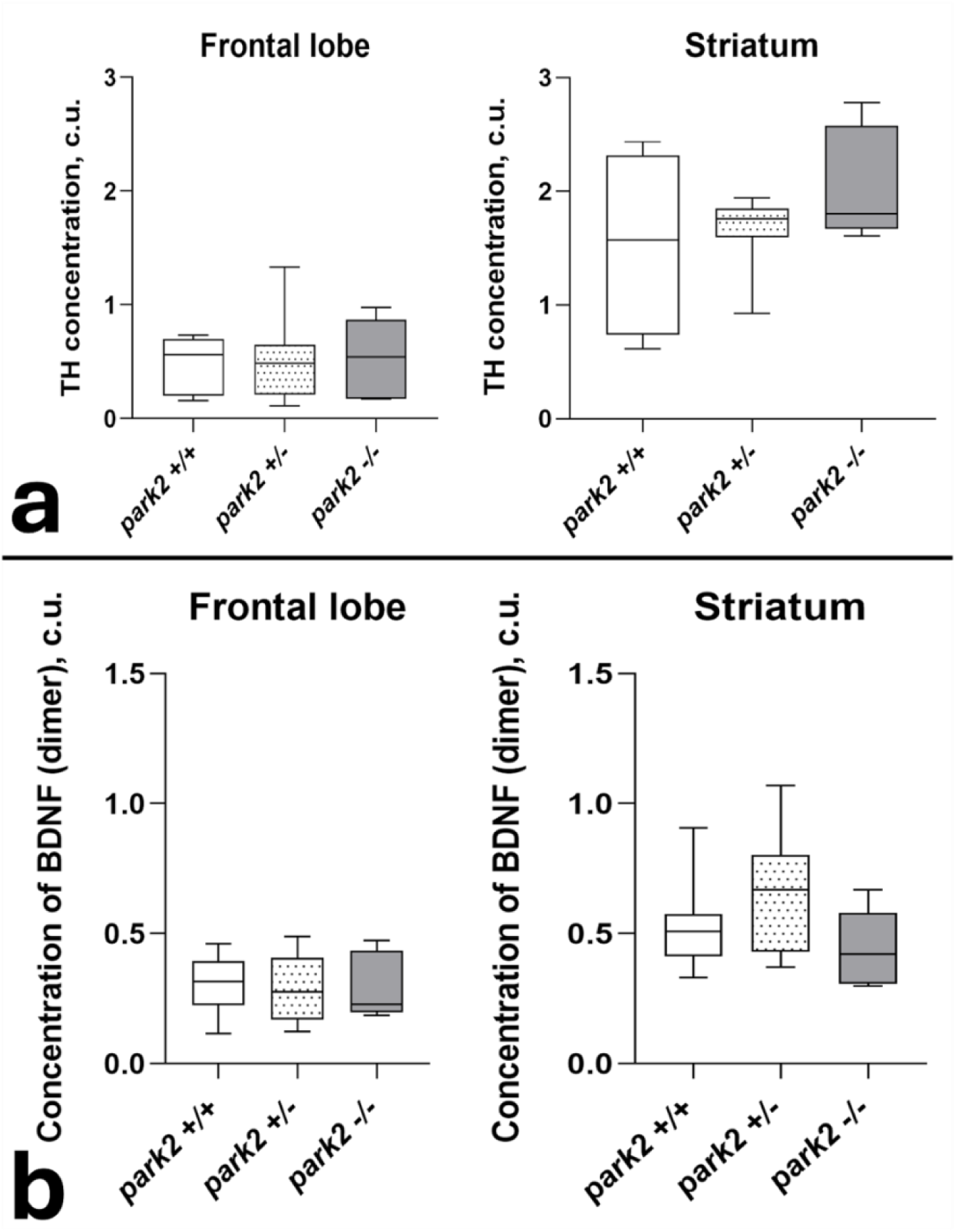
The content of **(a)** TH and **(b)** BDNF in the frontal lobe and striatum of 12-months male mice park2 +/+ (n=8), park2 +/- (n=8), and park2 -/- (n=5) measured by Western blotting. The data is presented as a median, the spread is shown as an interquartile range. One-way ANOVA, Dunn’s multiple comparisons test.

It is known that there is a significant decrease in the number of neurotrophic factors, such as BDNF and GDNF, with the progression of neurodegenerative diseases in the brain. Therefore, we measured their content in different brain structures using Western blotting.

BDNF performs the maturation of neurons, the formation of synapses, and has been shown to promote the survival of dopaminergic neurons. At the same time, its precursor proBDNF (32 kDa) acts through the p75 receptor and is responsible for the initiation of apoptosis, whereas monomeric (14 kDa) and dimeric forms (28 kDa) act as neurotrophic factors and promote cell survival.

We found that there were also no differences in the concentration of proBDNF, as well as its monomeric and dimeric forms in the striatum and in the frontal cortex in mice aged 12 months (Fig.3, b).

Another brain neurotrophin GDNF function as homodimer for tyrosine kinase receptors. The most studied feature of GDNF is its ability to support the survival of dopaminergic neurons and prevent chronic neurodegeneration. Therefore, its study in the context of PD is of particular interest. We detected 5 different isoforms of GDNF: (α)pro-GDNF, (β)pro-GDNF, mature GDNF, including two (34 and 32 kDa), which were not previously presented in the literature. Whether these proteins are additional isoforms of GDNF or represent products of its metabolism is currently unclear and future work is needed to expand our understanding of these results.

Assessment of GDNF content showed a decrease in the mature form of GDNF in knockout mice (Fig. 4). At the same time, there is an increase in the number of large-sized isoforms of GDNF, which probably represent inactive precursors of mature GDNF, which reflects a slowdown in the maturation rate of this neurotrophin. There is also an increase in (β)pro-GDNF in heterozygous mice in the frontal cortex. It is possible that the observed changes reflect the involvement of the Parkin protein in the processes of glial cell differentiation. The mechanisms and consequences of these changes have not been studied to date.

**Fig 4.**
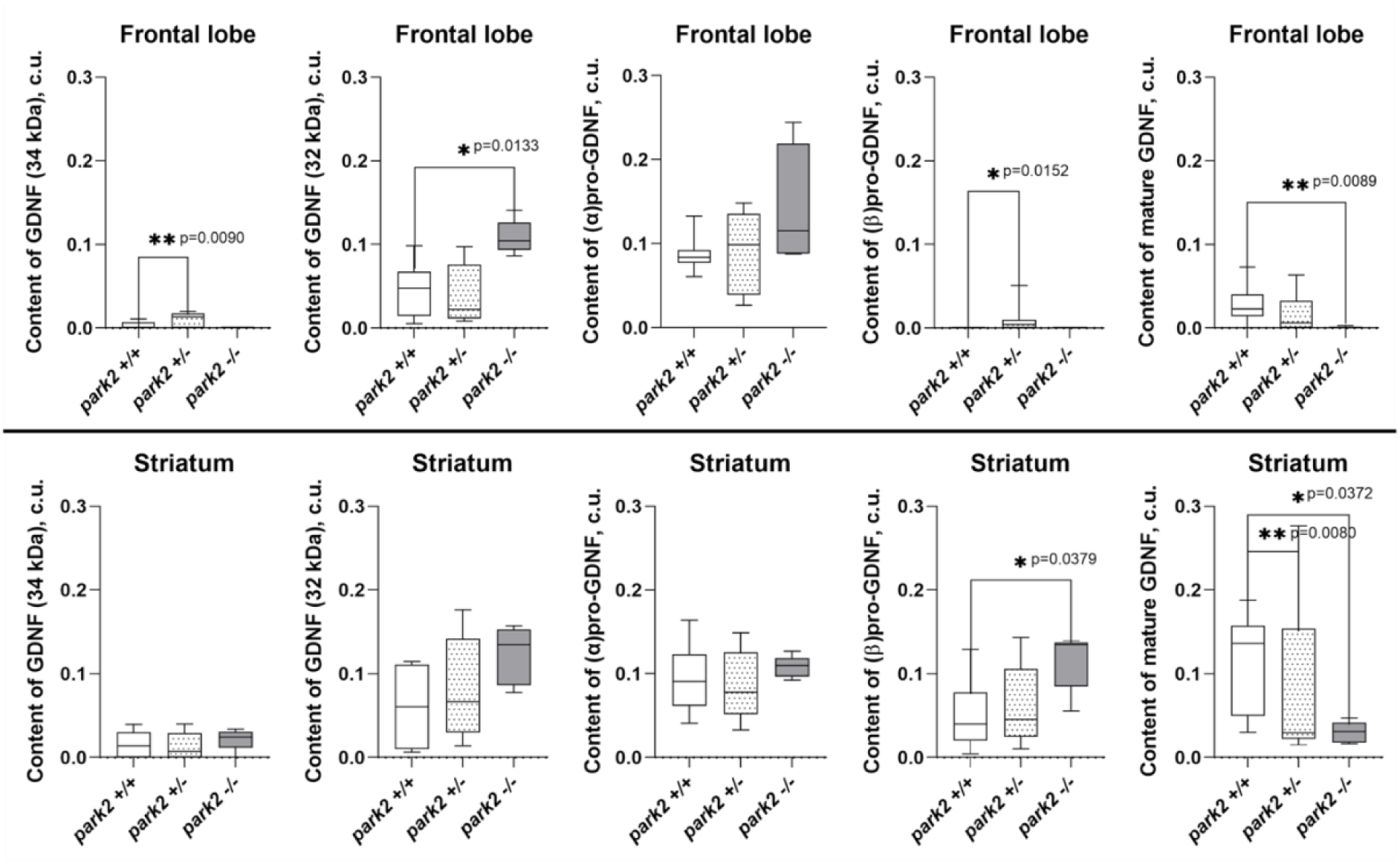
GDNF content in the frontal lobe and striatum of 12-months male mice park2 +/+ (n=8), park2 +/- (n=8), and park2 -/- (n=5) measured by Western blotting. The data is presented as a median, the spread is shown as an interquartile range. One-way ANOVA, Dunn’s multiple comparisons test.

## Conclusion

We studied the behavioral and biochemical parameters of mice knocked out by the *PARK2* gene. It was found that a distinctive feature of the behavior of these animals is a decrease in the level of anxiety and depression in adulthood with an earlier onset of motor deficiency (starting at about 16 months). However, the behavior of the control mice is practically compared to the experimental group at a later age. Among all the measured markers of PD (synthesis and metabolism of bioamines, as well as the amount of BDNF and GDNF) only a decrease in the level of mature GDNF in the brain of mice at the age of 12 months was observed; the rest did not differ from those in wild-type mice.

The data obtained indicates that mutations in the *PARK2* gene and, accordingly, a malfunction of the Parkin protein, although they lead to some neurodegenerative processes, are different from the symptoms of Parkinson’s disease in human. Moreover, if mutations in the *PARK2* gene in humans lead to the development of an early autosomal recessive form of the disease, then *park* -/- mice show some signs of decreased motor activity only by 12 months of life or more. It should be emphasized that most research in this area is limited to 1 year of animal life or less.^25^ The majority of them showed that Parkin’s genetic deletion causes disorders of the synaptic plasticity of the hippocampus, leading to memory deficits, without serious olfactory, emotional, or motor disorders, implying that the detected disorders will worsen with age and lead to the detailed symptoms of PD. The dynamics of behavioral and biochemical changes throughout the life of mice (up to 25 months) were studied in this research, which not only did not reveal an increase in specific signs of PD but showed a smoothing of differences. Therefore, the disruption of this gene in mice is not directly related to the development of parameters of Parkinson’s disease.

## Funding agencies

This study was supported by the advanced Russian ministry of science grant 075-15-2024-539.

## Financial Disclosures of all authors (for the preceding 12 months)

**1. Ekaterina A. Emelianova**

*Stock Ownership in medically-related fields* none

*Intellectual Property Rights* none

*Consultancies* none

*Expert Testimony* none

*Advisory Boards* none

*Employment* BostonGene Corporation (until 30 Aug 2024). Vardanants Str. 22, 0070, Yerevan, Armenia

*Partnerships* none

*Inventions* none

*Contracts* none

*Honoraria* none

*Royalties* none

*Patents* none

*Grants* Russian ministry of science grant 075-15-2024-539.

*Other* none

**2. Olga A. Averina**

*Stock Ownership in medically-related fields* none

*Intellectual Property Rights* none

*Consultancies* none *Expert Testimony* none *Advisory Boards* none *Employment* none *Partnerships* none *Inventions* none *Contracts* none *Honoraria* none *Royalties* none

*Patents* none

*Grants* Russian ministry of science grant 075-15-2024-539.

*Other* none

**3. Oleg A. Permyakov**

*Stock Ownership in medically-related fields* none

*Intellectual Property Rights* none

*Consultancies* none *Expert Testimony* none

*Advisory Boards* none

*Employment* none

*Partnerships* none *Inventions* none

*Contracts* none

*Honoraria* none

*Royalties* none

*Patents* none

*Grants* Russian ministry of science grant 075-15-2024-539.

*Other* none

**4. Anastasia V. Priymak**

*Stock Ownership in medically-related fields* none

*Intellectual Property Rights* none

*Consultancies* none

*Expert Testimony* none

*Advisory Boards* none

*Employment* none

*Partnerships* none

*Inventions* none

*Contracts* none

*Honoraria* none

*Royalties* none

*Patents* none

*Grants* Russian ministry of science grant 075-15-2024-539.

*Other* none

**5. Mariia A. Emelianova**

*Stock Ownership in medically-related fields* none

*Intellectual Property Rights* none

*Consultancies* none

*Expert Testimony* none

*Advisory Boards* none

*Employment* none

*Partnerships* none

*Inventions* none

*Contracts* none

*Honoraria* none

*Royalties* none

*Patents* none

*Grants* Russian ministry of science grant 075-15-2024-539.

*Other* none

**6. Olga O. Grigoryeva**

*Stock Ownership in medically-related fields* none

*Intellectual Property Rights* none

*Consultancies* none

*Expert Testimony* none

*Advisory Boards* none

*Employment* none

*Partnerships* none

*Inventions* none

*Contracts* none

*Honoraria* none

*Royalties* none

*Patents* none

*Grants* Russian ministry of science grant 075-15-2024-539.

*Other* none

**7. Svetlana A. Garmash**

*Stock Ownership in medically-related fields* none

*Intellectual Property Rights* none

*Consultancies* none

*Expert Testimony* none

*Advisory Boards* none

*Employment* none

*Partnerships* none

*Inventions* none

*Contracts* none

*Honoraria* none

*Royalties* none

*Patents* none

*Grants* Russian ministry of science grant 075-15-2024-539.

*Other* none

**8. Petr V. Sergiev**

*Stock Ownership in medically-related fields* none

*Intellectual Property Rights* none

*Consultancies* none

*Expert Testimony* none

*Advisory Boards* none

*Employment* none

*Partnerships* none

*Inventions* none

*Contracts* none

*Honoraria* none

*Royalties* none

*Patents* none

*Grants* Russian ministry of science grant 075-15-2024-539.

*Other* none

**9. Olga U. Frolova**

*Stock Ownership in medically-related fields* none

*Intellectual Property Rights* none

*Consultancies* none

*Expert Testimony* none

*Advisory Boards* none

*Employment* Institute of Mitoengineering of MSU, 119192 Moscow, Russia

*Partnerships* none *Inventions* none *Contracts* none

*Honoraria* none

*Royalties* none

*Patents* none

*Grants* Russian ministry of science grant 075-15-2024-539.

*Other* none

**10. Kirill E. Kianitsa**

*Stock Ownership in medically-related fields* none

*Intellectual Property Rights* none

*Consultancies* none

*Expert Testimony* none

*Advisory Boards* none

*Employment* Center of Molecular and Cellular Biology, Skolkovo Institute of Science and Technology, 143025 Skolkovo, Russia

*Partnerships* none

*Inventions* none

*Contracts* none

*Honoraria* none

*Royalties* none

*Patents* none

*Grants* Russian ministry of science grant 075-15-2024-539.

*Other* none

**11. Vladislav S. Savitskiy**

*Stock Ownership in medically-related fields* none

*Intellectual Property Rights* none

*Consultancies* none

*Expert Testimony* none

*Advisory Boards* none

*Employment* Institute of Mitoengineering of MSU, 119192 Moscow, Russia

*Partnerships* none

*Inventions* none

*Contracts* none

*Honoraria* none

*Royalties* none

*Patents* none

*Grants* Russian ministry of science grant 075-15-2024-539.

*Other* none

**12. Maxim L. Lovat**

*Stock Ownership in medically-related fields* none

*Intellectual Property Rights* none

*Consultancies* none

*Expert Testimony* none

*Advisory Boards* none

*Employment* Institute of Mitoengineering of MSU, 119192 Moscow, Russia Biology Faculty, Lomonosov Moscow State University, 119991 Moscow, Russia *Partnerships* none

*Inventions* none

*Contracts* none

*Honoraria* none

*Royalties* none

*Patents* none

*Grants* Russian ministry of science grant 075-15-2024-539.

*Other* Teaching biology as part of additional education (working with schoolchildren).

